# simpleaf: A simple, flexible, and scalable framework for single-cell transcriptomics data processing using alevin-fry

**DOI:** 10.1101/2023.03.28.534653

**Authors:** Dongze He, Rob Patro

## Abstract

**Summary:** The `alevin-fry` ecosystem provides a robust and growing suite of programs for single-cell data processing. However, as new single-cell technologies are introduced, as the community continues to adjust best practices for data processing, and as the `alevin-fry` ecosystem itself expands and grows, it is becoming increasingly important to manage the complexity of `alevin-fry`’s single-cell preprocessing workflows while retaining the performance and flexibility that make these tools enticing. We introduce `simpleaf`, a program that simplifies the processing of single-cell data using tools from the `alevin-fry` ecosystem, and adds new functionality and capabilities, while retaining the flexibility and performance of the underlying tools.

**Availability and implementation:** `Simpleaf` is written in Rust and released under a BSD 3-Clause license. It is freely available from its GitHub repository https://github.com/COMBINE-lab/simpleaf, and via bioconda. Documentation for `simpleaf` is available at https://simpleaf.readthedocs.io/en/latest/ and tutorials for `simpleaf` are being developed that can be accessed at https://combine-lab.github.io/alevin-fry-tutorials.

## Introduction

Single-cell sequencing has become an indispensable tool for studying cellular biology at the resolution of individual cells [1, 2], and processing the resulting data often requires a dedicated suite of tools and methods. Recently, He et al. [3] demonstrated that the `alevin-fry` ecosystem provides an efficient, accurate, and flexible framework for single-cell data processing. Yet, the rapid arrival of new technologies and experimental modalities have led to data analysis pipelines that require increasingly complex and sophisticated workflows. For example, analyzing CITE-seq [4] data involves executing the entire `alevin-fry` pipeline three times, each time with a slightly different configuration and on different sets of files. Likewise, as improved tools, like the `piscem` [5, 6] read mapping tool, are introduced into the `alevin-fry` ecosystem, users wishing to adopt these new tools have to learn their interfaces and logistics.

### Software Description

To simplify and ease the user experience for both simple and complex experimental setups, and to allow seamless use of the newest `alevin-fry` ecosystem components, we have developed `simpleaf` (simple `alevin-fry`). `Simpleaf` (overview in Figure 1) is a high-level framework, that provides simple, flexible, and scalable interfaces for uniformly accessing standard and advanced features in the `alevin-fry` ecosystem.

**Fig. 1.**
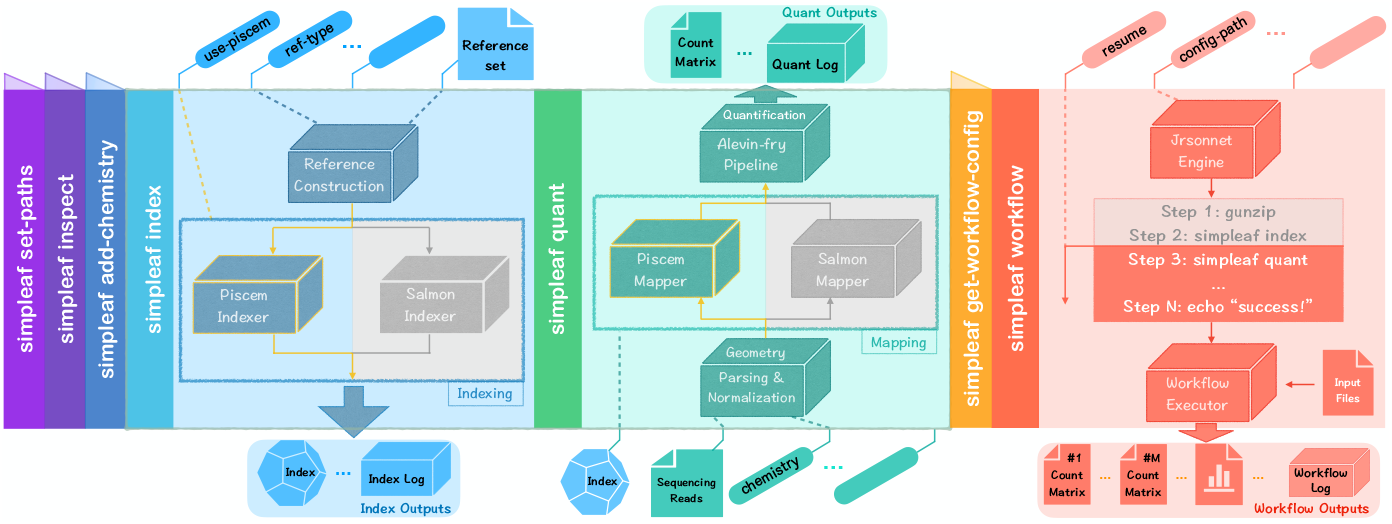
Overview of some salient `simpleaf` subcommands, showing the flow of data through a hypothetical invocation. The leftmost expanded column (blue) represents using the `simpleaf index` command to build a reference sequence and the corresponding `piscem` index. The center expanded column (aqua) represents using this `piscem` index in conjunction with sequenced reads to produce a count matrix for subsequent analysis. Note that the indexer and mapper of both `piscem` and `salmon` are fully supported in `simpleaf`. Finally, the rightmost expanded column (red) represents the invocation of a hypothetical `simpleaf workflow`, where the workflow can require several input files and produce several outputs. Additional `simpleaf` subcommands are described in the appendix.

The concept of “wrapper” or “workflow” programs in the context of single-cell data processing pipelines is well-established. Apart from the many bespoke workflows developed for individual technologies, there exist several tools designed to ease and simplify the processing of multiple types of data. Here, we highlight a few examples, though this is not intended, to constitute an exhaustive list of such tools. The popular `Cell Ranger` [7] tool itself is, in part, a Python script that wraps `STAR` [8] and other tools designed by 10x Genomics. The `zUMIs` [9] and `UniverSC` [10] tools provide highly-capable suites of workflows for processing data generated using many different technologies using, respectively, `STAR` and `Cell Ranger` itself. `Kb-python` is a Python tool that wraps `kallisto|bustools` [11] and related tools, and provides a high-level interface to process data from many different experimental protocols and setups. The `scPipe` [12] tool is a modular wrapper around multiple tools and packages within the R ecosystem, which uses the Subread aligner [13] for mapping, and is capable of processing data from a variety of different single-cell technologies.

`Simpleaf` is dedicated to providing a simple and flexible user interface for the `alevin-fry` ecosystem, which consists of a range of underlying tools and modules for single-cell data processing [3, 5, 6, 14, 15, 16]. In addition to coordinating the execution of these tools and providing a unified and simplified interface, `simpleaf` also adds new functionality aimed at further generalizing the capabilities of the underlying tools, and allowing users to easily create and share their own workflows without having to program or modify `simpleaf` itself.

### A simplified interface to core alevin-fry functionality

The most basic functionality provided by `simpleaf` is that it provides a simplified yet flexible interface to the underlying `alevin-fry` modules and capabilities.

In `simpleaf`, the standard `alevin-fry` pipeline [3] is distributed over two phases: (i) indexing, which includes creating an augmented (*splici* [3] or *spliceu* [6]) reference, where appropriate, and building the corresponding reference index, and (ii) quantification, which consists of read mapping, cell barcode detection and correction, and UMI resolution. These two phases are exposed as two sub-commands within `simpleaf`: `simpleaf index` and `simpleaf quant`. Each of these, in turn, exposes various flags for retaining critical flexibility in processing.

Although most of the functionality provided by `simpleaf` programs can be directly replicated by calling the underlying tools with the appropriate configurations and arguments, the advantage of using `simpleaf` comes from the fact that `simpleaf`, by default, incorporates the best practices for running the underlying tools, reduces the workload by automatically handling tedious but crucial details one needs to take care of in the most common use cases, and also retains critical flexibility when necessary. For example, if a `simpleaf index` invocation is followed by a call to `simpleaf quant`, `simpleaf quant` will automatically recruit and parameterize the correct mapper, and will automatically locate and provide the file containing the transcript-to-gene mapping information to later quantification stages where appropriate. This file would have explicitly provided if `alevin-fry` is not invoked through `simpleaf`. Yet, to provide for maximum flexibility, `simpleaf` provides alternative processing options as well, like the option to begin the quantification process from an already-computed set of mapping results and thus to skip the mapping process.

### Dedicated parameterization for easily switching between options and underlying tools

As the methodologies underlying single-cell quantification advance, we have continued to improve the existing features of the `alevin-fry` ecosystem and introduce new options and functionality wherever appropriate. Yet, the options and possibilities for single-cell analysis continue to expand, and the burden on users to keep up with new methods, tools, and best practices grows. To ensure that the best practices and new features of the `alevin-fry` ecosystem can be easily accessed and applied by users to their data, `simpleaf` tracks current best practices and provides built-in parameterizations under simplified configurations, which allows seamless switching between different options and backend tools.

One example is the ability to easily switch between `piscem` [5, 6] and `salmon` [16, 14] as the underlying reference indexing and mapping tools. `Piscem` is a new index and mapper in the `alevin-fry` ecosystem that further lowers the memory requirements for single-cell data processing [6], and `salmon` [16, 14], which relies on the `pufferfish` index [17], is the traditional mapper used with `alevin-fry`. As `piscem` and `salmon` are independent tools with distinct parameters, explicitly switching from `salmon` to `piscem` requires knowledge about the `piscem` tool and the relevant details of its indexing and mapping subcommands. However, in `simpleaf`, the only modification needed to make use of `piscem` is to pass the `--use-piscem` flag. Furthermore, for simplicity, if this flag is set when calling `simpleaf index`, the subsequent `simpleaf quant` executions that map against this index will automatically use `piscem`, with appropriate parameters, as the mapper.

Another example is the ability to seamlessly build different types of reference indices by simply changing the flags passed to the `simpleaf index` command. Currently, `simpleaf` has the ability to build three kinds of reference indices. Although the procedure for generating these types of reference indexes is different, `simpleaf` abstracts over the technicalities and only requires the user to set the `--ref-type` option as desired.

### Parsing protocols with a complex fragment geometry

Another useful feature provided by `simpleaf` is the ability to represent and parse fragment geometry specifications that are potentially *more complex* than those directly supported by the underlying mappers — for example, those with variable barcode length and floating barcode position, such as sci-RNA-seq3 [18] — using a concise description language^1^. An example of streaming parser invocation can be found in appendix section B. When presented with a “complex” geometry specification, `simpleaf` “normalizes” these reads into an appropriate “simple” format (a format with only known position and fixed-length barcodes and UMIs) on the fly, and provides a modified (and simplified) geometry format description to the underlying mapper. Moreover, it streams the normalized reads directly to the mapper using FIFOs programmatically managed by `simpleaf`, thereby avoiding intermediate disk usage. This feature enables the existing mappers in the `alevin-fry` ecosystem, which are designed to process reads with simple geometry, to handle sophisticated geometries without modifying the underlying mapping tools, requiring extra preprocessing from the users, or taking the extra time and space to write and read the intermediate representations.

To date, the community has put significant efforts into documenting and categorizing the library layout for many existing sequencing assays [19, 10, 20] and developing general parsers for such protocols [21, 22, 10, 23]. Of course, these tools, or their relevant components could also be applied to this task, with the user handling the appropriate bookkeeping. However, the built-in capability of `simpleaf` focuses on providing a concise language for representing both simple and complex fragment geometry that can be passed directly to `simpleaf` from the command line, and the seamless internal normalization of this complex geometry into a simplified form compatible with both supported mappers.

### Generalized and sharable workflow construction for complex single-cell workflows

`Simpleaf` also provides the ability to execute complex and highly-configurable `alevin-fry` workflows described by simple user-provided configuration files. The purpose of the `simpleaf` workflow module is neither to replace general workflow languages like Nextflow [24] or Snakemake [25] that enable near-limitless generality, but that require learning sophisticated and complex domain-specific languages, nor to expose some set of easy-to-use but pre-defined workflows for complex single-cell protocols as in `Cell Ranger` and `kb-python`. Rather, `simpleaf workflow` aims to provide a platform for `alevin-fry` users with all levels of programming knowledge to easily create, invoke, and share their workflows, which can contain not only `simpleaf` commands but also any shell commands that are valid in the user’s terminal. It allows the definition, via a simple imperative configuration and templating system, of custom workflows parameterized on user-defined input, which can then be reused to simplify the processing of complex workflows and easily shared with other users.

`Simpleaf workflow` takes a workflow configuration (a Jsonnet program) as input^2^, converts it to a complete workflow JSON file, and invokes some or all recorded `simpleaf` program commands and external shell commands in an appropriate order. More details can be found in Appendix sections A.7 and C. This design allows users with limited programming experience to define a simple but useful workflow as a JSON configuration, but also makes it possible for advanced users to develop sophisticated Jsonnet programs to generate `simpleaf` workflows by taking advantage of the full functionality provided by the Jsonnet language. Furthermore, this design also makes it easy for users to create and share their workflows by simply sharing their workflow configuration files, without the need to understand and contribute to the codebase of `simpleaf` itself. To demonstrate the utility of the `simpleaf workflow` command, we have built such workflow configurations for processing CITE-seq [4] and 10x Genomics feature barcode data. We are continuing to develop more workflow configurations and are also accepting contributions from the community.

## Discussion

`Simpleaf` provides a simple and flexible interface to access the state-of-the-art features provided by the `alevin-fry` ecosystem, tracks best practices using the underlying tools, enables users to transparently process data with complex fragment geometry, and to build and execute sophisticated workflows containing both `simpleaf` and external commands without the need to write code. `Simpleaf` has already seen adoption in community-led projects, for example, in the `scrna` analysis pipeline of the nf-core project [26, 27]. We hope that, in the future, `simpleaf` can serve as an entry point and main interface to the `alevin-fry` ecosystem for most users. We also envision that our workflow feature can encourage those in the community to create and share their workflows and, thus, can help `simpleaf` to provide increasingly reusable building blocks to enable more varied and sophisticated single-cell data analysis pipelines.

While `simpleaf` provides a simple and flexible framework for single-cell data processing, the current implementation still has some limitations, which motivate future work. For example, although the fragment geometry parser can parse barcodes with variable length and floating position, it currently lacks the ability to perform certain kinds of preprocessing, like the barcode substitution scheme required by the split-seq [28] technology. This can be solved by expanding the current geometry specification to describe and enable this kind of preprocessing. Moreover, `simpleaf workflow` is still under active development. Current efforts are underway to improve its generality, expand its library of standard functions, and to develop more useful and sophisticated workflows for different purposes.

## Competing interests

RP is a co-founder of Ocean Genomics inc.

## Funding

This work has been supported by the US National Institutes of Health (R01 HG009937), and the US National Science Foundation (CCF-1750472, and CNS-1763680). Also, this project has been made possible in part by grant number 252586 from the Chan Zuckerberg Initiative Foundation. The funders had no role in the design of the method, data analysis, decision to publish or preparation of the manuscript.

## Appendices

### Appendix A simpleaf subcommands

`Simpleaf` provides subcommands for various purposes. Here we briefly discuss the current subcommands exposed in `simpleaf`. The `alevin-fry` team is actively working to improve existing modules in the `alevin-fry` ecosystem as well as developing new modules that will be most useful for the community. For complete and up-to-date documentation, one can refer to the official `simpleaf` documentation at https://simpleaf.readthedocs.io/en/latest/.

In order to operate properly, `simpleaf` requires that the environment variable `ALEVIN FRY HOME` exists. It will use the directory pointed to by this variable to cache useful information (e.g. the paths to selected versions of the tools it invokes, configurations containing the mappings for custom chemistries that have been registered, and other information like the permit lists for certain chemistries). In popular shells such as `bash` and `zsh`, this variable be set with `export ALEVIN FRY HOME=/full/path/to/dir/you/want/to/use`

#### A.1 simpleaf set-paths

Before calling `simpleaf`, one has to run the `simpleaf set-paths` command to set the paths to the underlying tools. If no flags are provided, this program will try to find the dependencies in the shell’s `PATH`. Currently, to use `simpleaf`, one must provide a compatible version of `pyroe` (https://pyroe.readthedocs.io/en/latest/), `alevin-fry` (https://alevin-fry.readthedocs.io/en/latest/), and at least one of the `alevin-fry` mappers, `piscem` or `salmon`.

#### A.2 simpleaf inspect

This subcommand inspects the configuration of `simpleaf` in the current environment — such as the path to its dependencies and the custom chemistries that have been registered — and reports on the current configuration.

#### A.3 simpleaf index

The `simpleaf index` subcommand provides the functionality to build a reference index from the provided reference set using either the default `salmon` or the improved `piscem` indexing tool. Usually, for a species, protocol pair, `simpleaf index` needs only to be run once, and all subsequent experiments can utilize that index. By default, `simpleaf index` takes a reference genome in FASTA format and gene annotation in GTF format, and makes the *spliced+intronic* (*splici*) reference index after extracting the sequence of the spliced transcripts and intronic regions. Other augmented reference types, such as the *spliced+unspliced* (*spliceu*) reference, are also available in `simpleaf index` by setting the `--ref-type` argument appropriately.

If one would like to build the index directly from the provided reference sequences, for example, when the genome build of a species is unavailable and the transcriptome sequences are provided directly, one can pass the reference sequence file to the `--ref-seq` argument.

#### A.4 simpleaf quant

The `simpleaf quant` subcommand is designed as an all-in-one program to generate a gene*×*barcode count matrix directly from the provided index and sequencing reads. By default, it takes a `piscem` or `salmon` index, and the sequencing read (lists of `FASTQ` files) as input. It maps the reads and resolves the associated UMIs after detecting and correcting the cellular barcodes. It also provides the option to directly begin quantification from an already-computed set of mapping results, thereby skipping the mapping process, via the `--map-dir` argument.

#### A.5 simpleaf add-chemistry

This subcommand adds a new custom fragment geometry specification to the `simpleaf` geometry library together with a unique name. Once added, one can specify that custom geometry specification using the associated name when running `simpleaf quant`. For example, if one wants to store the geometry specification of sci-RNA-seq3, one can invoke `simpleaf add-chemistry --name sci-seq3 --geometry` “`1`*{*b[9-10]f[CAGAGC]u[8]b[10]*}*`2`*{*`r:`*}*”. Then, one can use this geometry when calling `simpleaf quant` by specifying `--chemistry sci-seq3`.

#### A.6 simpleaf get-workflow-config

This subcommand gets the workflow configuration file of a published workflow from the protocol estuary GitHub repository (https://github.com/COMBINE-lab/protocol-estuary). When calling this program, `simpleaf` will automatically download the protocol estuary GitHub repository into the `ALEVIN FRY HOME` directory. If invoking unpublished workflows, one can skip this step and provide the workflow configuration files directly to `simpleaf workflow` via the `--config-file` flag.

#### A.7 simpleaf workflow

The `simpleaf workflow` subcommand is designed to run potentially complex single-cell data processing workflows according to a `simpleaf` workflow configuration file. Any valid Jsonnet ((https://jsonnet.org/)) program and JSON file is a valid simpleaf workflow configuration file. In `simpleaf workflow`, the provided configuration file will be first converted to a `simpleaf` workflow JSON file. Whereas the Jsonnet file provides a “template” for the workflow and functions to handle features like basic logic, the workflow JSON file that results is a simple imperative description of the commands that are to be executed.

To ease the later parsing process, the values of all command arguments in the configuration file must be provided as strings, i.e., wrapped by quotes (“`value`”). `Simpleaf workflow` will traverse the converted workflow JSON file to find and parse the valid fields that record either a `simpleaf` or an external shell command. To provide the greatest flexibility, the only requirement `simpleaf workflow` sets is for the layout of the fields that records a command, either a `simpleaf` command or an external command.

1. A command record field must contain a `Step` and a `Program Name` sub-field, where the `Step` field represents which step, using an integer, this command constitutes in the workflow. The `Program Name` field represents a valid program in the user’s execution environment. For example, the correct `Program Name` for `simpleaf index` is “`simpleaf index`”. For an external command such as `awk`, if its binary is in the user’s `PATH` environmental variable, it can just be “`awk`”; if not, it must contain a valid path to its binary, for example, “`/usr/bin/awk`”.
2. If a field records a `simpleaf` command, i.e., it has a valid `Step` and `Program Name` field, the name of the rest of its sub-fields must be valid `simpleaf` flags (for example, options like `--fasta`, or `-f` for short, for `simpleaf index` and z`--unfiltered-pl`, or `-u` for short for `simpleaf quant`). Those option names (sub-field names), together with their values,if any, will be used to call the corresponding `simpleaf` program. Sub-fields that are not named by a valid `simpleaf` flag will be ignored.
3. If a field records an external shell command, it must contain a `Step` and a `Program Name` sub-field as described above. In contrast to `simpleaf` command records, all arguments of an external shell command must be provided in an array, in order, with the name “`Arguments`”. `Simpleaf workflow` will parse the entries in the array to build the actual command. For example, to tell `simpleaf workflow` to invoke the shell command $`ls -l -h.` at step 7, one needs to use the following JSON record:

~~~
{
 “Step”: 7,
 “Program Name”: “ls”,
 “Arguments”: [“-l”, “-h”, “.”]
}
~~~

After converting the workflow configuration file into a `simpleaf` workflow JSON file, `simpleaf workflow` will parse and invoke the commands in the workflow according to the following flags. If none of the flags are set, `simpleaf workflow` will invoke the complete workflow.

- If setting the `no-execution` flag, `simpleaf workflow` will write the complete workflow JSON file and other log files but invoke none of the commands.
- If setting the `start-at` flag with a step number, `simpleaf workflow` will begin the invocation from that specific step number according to the `Step` field in the command records.
- If setting the `resume` flag, `simpleaf workflow` will find the JSON configuration of a previous `simpleaf workflow` run in the provided output folder to decide which is the starting step number.
- If setting the `skip-step` flag with a set of comma-separated step numbers, `simpleaf workflow` will invoke all but the commands represented by those skipped step numbers.

Many useful and frequently used functions have been provided as a `simpleaf` workflow utility library at https://github.com/COMBINE-lab/protocol-estuary, which is also the place we recommend our users submit the workflows they designed.

### Appendix B Example of streaming parser invocation

Internally, `simpleaf` will first check if the given geometry specification has a fixed barcode length and position. If it does, read files will be directly passed to the underlying mapper, designed to take fixed geometry reads. If not, `simpleaf` will pass the geometry specification to its sequence geometry parser (https://github.com/COMBINE-lab/seq_geom_parser) and call its sequence geometry transformer (https://github.com/COMBINE-lab/seq_geom_xform) to “normalize” the reads to a fixed geometry format and stream the transformed reads directly to the mapper, without writing intermediate files. For example, the simplified geometry description for the geometry originally specified in Appendix section A.5 is `1`*{*b[11]u[8]b[10]*}*`2`*{*`r:`*}*. The length of the first barcode changes from 9 or 10 nucleotides to 11 because the sequence geometry transformer augments the first barcode segment with an `A` if the original barcode is of length 10 or with `AC` if it is of length 9. By doing so, the first barcode of all reads will be of the same length, and the augmented barcodes of hairpin barcodes of different lengths will not overlap, as all hairpin barcodes of length 9 now end with `C` and all of the length 10 ends with `A` (this scheme can be naturally generalized to different segment width ranges).

### Appendix C Example of simpleaf workflow invocation

As discussed in section A.7, `simpleaf workflow` is designed to generate and invoke single-cell analysis workflows according to a workflow configuration file (Jsonnet program). `Simpleaf` uses the Jrsonnet (https://github.com/CertainLach/jrsonnet) library for parsing the underlying Jsonnet configuration.

In the protocol estuary GitHub repository (https://github.com/COMBINE-lab/protocol-estuary), we have published a workflow for processing CITE-seq data. Once one has obtained the workflow configuration file, for example, by calling `simpleaf get-workflow-config` with the argument `--workflow cite-seq 10xv2` as discussed in Appendix section A.6 and has filled in all required fields in the configuration file, i.e., the file path to the needed files, they can call `simpleaf workflow` and pass the complete configuration file to the `--config-file` argument. `simpleaf workflow` will convert the configuration file to a workflow JSON file using Jrsonnet and recursively find and parse all valid command records in the JSON file.

Without the help of `simpleaf workflow`, one needs to develop and invoke 12 distinct commands in the shell, including preprocessing, to obtain all the desired results of this specific workflow. Using `simpleaf workflow`, one only needs to fill the path to the files of sequencing reads and reference sets in the configuration file, and pass it to `simpleaf workflow` via the `--config-file` argument. Then, `simpleaf workflow` will automatically generate a data analysis pipeline for that specific dataset and invoke all required commands for analyzing the data.

## Footnotes

1 The current specification of this description language is available at https://hackmd.io/@PI7Og0l1ReeBZu_pjQGUQQ/rJMgmvr13.

2 Any valid JSON document is also a valid Jsonnet program.

## Notes

https://github.com/COMBINE-lab/simpleaf

